# Eye movements decrease during effortful speech listening

**DOI:** 10.1101/2023.02.08.527708

**Authors:** M. Eric Cui, Björn Herrmann

**Affiliations:** Rotman Research Institute, Baycrest, M6A 2E1, North York, ON, Canada; Department of Psychology, University of Toronto, M5S 1A1, Toronto, ON, Canada

**Keywords:** Eye-tracking, listening effort, eye movements, pupillometry, speech processing, spoken stories

## Abstract

Pupillometry is the most used objective tool to assess listening effort but has several disadvantages. The current study explores a new, objective way to assess listening effort through eye movements. Building on cognitive and neurophysiological work, we examine the hypothesis that eye movements decrease when speech listening becomes challenging. In three experiments with human participants from both sexes, we demonstrate, consistent with this hypothesis, that fixation duration increases and spatial gaze dispersion decreases with increasing speech masking. Eye movements decreased during effortful speech listening for different visual scenes (free viewing; object tracking) and speech materials (simple sentences; naturalistic stories). In contrast, pupillometry was insensitive to speech masking during story listening, highlighting the challenges with pupillometric measures for the assessments of listening effort in naturalistic speech-listening paradigms. Our results reveal a critical link between eye movements and cognitive load, and provide the foundation for a novel measure of listening effort applicable in a wide range of contexts.

**Significance statement:** Assessment of listening effort is critical for early diagnosis of age-related hearing loss. Pupillometry is most used but has several disadvantages. The current study explores a new, objective way to assess listening effort through eye movements. We examine the hypothesis that eye movements decrease when speech listening becomes effortful. We demonstrate, consistent with this hypothesis, that fixation duration increases and gaze dispersion decreases with increasing speech masking. Eye movements decreased during effortful speech listening for different visual scenes (free viewing; object tracking) and speech materials (sentences; naturalistic stories). Our results reveal a critical link between eye movements and cognitive load, and provide the foundation for a novel measure of listening effort applicable in a wide range of contexts.

## Introduction

Hearing impairment affects about 40% of people aged over 60 years (Feder et al., 2015; Goman and Lin, 2016), but is often diagnosed decades after speech-comprehension difficulties in noisy situations, such as crowded restaurants, emerge (Pichora-Fuller et al., 1995; Pichora-Fuller and Levitt, 2012). Individuals with even mild hearing impairment rely substantially on attention in noisy situations, which makes listening effortful (Pichora-Fuller et al., 2016; Peelle, 2018; Herrmann and Johnsrude, 2020b). Accurate assessment of listening effort may thus help diagnose hearing impairment earlier and evaluate treatment outcomes. However, existing measures have limitations that hinder their use in practice. The current study aims to explore a novel approach to assess speech-comprehension difficulties associated with listening effort.

Self-reports via subjective ratings are a common way to assess listening effort (Gatehouse and Noble, 2004; Larsby et al., 2005; Krueger et al., 2017). However, subjective measures can be influenced by different reference frames and attribution effects (Moore and Picou, 2018). Physiological measures provide an objective window onto listening effort that can remedy disadvantages of subjective ratings (Mackersie et al., 2015; Wöstmann et al., 2015; Dimitrijevic et al., 2017; Miles et al., 2017). Pupillometry – the measurement of pupil dilation – is the most used objective tool to assess listening effort (Winn et al., 2018; Zekveld et al., 2018; Kadem et al., 2020). However, pupil dilation is sensitive to environmental changes in luminance (Knapen et al., 2016) and light spectrum (Suzuki et al., 2019; Thurman et al., 2021), and is therefore difficult to use outside of the laboratory. Moreover, measuring pupil dilation accurately requires participants to continuously fixate on a point on a computer monitor (Ohlenforst et al., 2017; Zekveld et al., 2018; Farahani et al., 2020; Winn and Teece, 2021), because luminance changes arising from eye movements change the pupil dilation and different angles of the pupil relative to the eye-tracker can make the pupil diameter appear different without an actual difference (Brisson et al., 2013; Hayes and Petrov, 2016; Fink et al., 2021). Yet, restriction of gaze to a central fixation point creates a dual task that is uncommon in everyday life and central fixation can impair memory and mental imagery for spoken speech (Johansson et al., 2012).

A few attempts have been made to assess listening effort using the small, jerk-like, involuntary eye movements during fixation (microsaccades) that can be measured concurrently with pupillometry (Kadem et al., 2020). However, microsaccades appear neither sensitive to acoustically-nor linguistically-induced listening effort (Kadem et al., 2020). Microsaccade amplitudes are also very small (Martinez-Conde et al., 2009; Martinez-Conde et al., 2013), requiring high-resolution recording (Poletti and Rucci, 2016), and microsaccade recordings suffer from the same disadvantages associated with forced gaze fixation as does pupillometry. These limitations hinder the utility of pupillometric and microsaccadic indices for the assessment of speech-comprehension difficulties in noise and associated listening effort.

Individuals naturally explore their environments through eye movements (Fukushima, 2003; Ono, 2015; Missal and Heinen, 2017) and visual exploration can be incidental while engaged in a different task (Lipton et al., 1980; Hutton and Tegally, 2005; Kosch et al., 2018). Such incidental eye movements may provide a window onto cognition, since people avert gaze (Glenberg et al., 1998), reduce object-tracking eye movements (Lipton et al., 1980; Hutton and Tegally, 2005; Kosch et al., 2018), and decrease saccades (Walter and Bex, 2021) during periods of high memory load, raising the possibility that all cognitively demanding tasks, including speech comprehension, affect eye movements. This possibility is supported by neurophysiological evidence showing that a reduction in movements increases neural activity in the auditory cortex and, in turn, improves sound perception (Schneider et al., 2014; McGinley et al., 2015; Schneider and Mooney, 2015, 2018). Critically, eye movements directly modulate neuronal excitability in auditory cortex, such that the likelihood of a neuron firing, and thus responding to sound, increases in the absence of eye movements (O’Connell et al., 2020). Research further suggests that eve movements and pupil dilation might be driven by common underlying neurophysiology (Joshi and Gold, 2020; Wang and Munoz, 2021; Burlingham et al., 2022), and, as a result, may perhaps both be sensitive to listening effort.

In the current study, we propose that leveraging eye movements to make inferences about audition will deliver an effective measure of listening effort. We suggest that when listening becomes effortful, eye movements decrease to free resources for speech comprehension. In three experiments with different speech materials and different visual-stimulation displays, we examine the hypothesis that eye movements are sensitive to speech masking that is associated with listening effort.

## General Methods

### Participants

Younger adults aged 18–34 years participated in the three experiments of the current study. Participants were either native English speakers or highly proficient non-native English speakers. Demographic information for each participant is provided in the sections describing the methods for each experiment. Participants gave written informed consent prior to the experiment and were paid $7.5 CAD per half-hour for their participation. Participants self-reported having normal hearing abilities. The study was conducted in accordance with the Declaration of Helsinki, the Canadian Tri-Council Policy Statement on Ethical Conduct for Research Involving Humans (TCPS2-2014), and was approved by the Research Ethics Board of the Rotman Research Institute.

### Stimulation and recording setup

Sounds were presented via Sony Dynamic Stereo MDR-7506 headphones and a Steinberg UR22 mkII (Steinberg Media Technologies) external sound card. Stimulation was run using Psychtoolbox (v3.0.14) in MATLAB (MathWorks Inc.) on a Lenovo T450s laptop with Microsoft Windows XP. The laptop screen was mirrored to an ASUS monitor with a refresh rate of 60 Hz. All sounds were presented at a comfortable listening level.

During the experiments, participants rested their head on a chin and forehead rest facing the computer monitor at a distance of about 70 cm. Pupil area and eye movements were recorded continuously from the right eye (or the left eye if the right eye could not be tracked accurately) using an integrated infrared camera (EyeLink 100 plus eye tracker; SR Research Ltd.) at a sampling rate of 500 Hz. Nine-point fixation was used for eye-tracker calibration prior to each stimulation block (McIntire et al., 2014).

### Preprocessing of eye-movement and pupil-area data

Preprocessing of eye-movement and pupil-area data involved removing eye blinks and other artifacts. For each eye blink indicated by the eye tracker, all data points between 100 ms before and 200 ms after a blink were set to NaN (‘not a number’ in MATLAB). In addition, pupil-area values that differed from the mean pupil area by more than 3 times the standard deviation were classified as outliers and set to NaN. Missing pupil data (coded as NaN) resulting from artifact rejections and outlier removal were interpolated using MATLAB’s ‘pchip’ method. X- and y-time courses were not interpolated (except for microsaccade/saccade analyses), but missing data points (NaNs) were ignored in eye-movement data-analysis procedures.

### Pupil and eye-movement metrics

Pupil-area time courses were filtered with a 5-Hz low-pass filter (51 points, Kaiser window). Because participants were not required to fixate in one location on the screen for most of the experimental conditions, pupil area could be affected by the changing angle of the pupil relative to the eye-tracking camera. In order to mitigate this potential issue, we regressed out any linear and quadratic relationship of x and y with the pupil area before low-pass filtering (Fink et al., 2021; Kinley and Levy, 2021; Kraus et al., 2022). There were no meaningful differences in the results reported below compared to the uncorrected pupil area.

Two main metrics were used to investigate whether eye movements changed depending on the degree to which speech was masked by background babble: fixation duration and spatial gaze dispersion. Fixation duration was calculated as the time a person’s x-y eye coordinates remained in a given location (within 0.5° visual angle; radius of 10 px). For each time point, the corresponding x-y coordinate defined the critical 0.5° visual-angle location. The number of continuous pre- and post-samples was calculated for which the x-y-eye coordinates remained in the critical location. The sample number was divided by the sampling frequency to obtain the fixation duration for the specific time point. If a data value of any pre- or post-sample within the 0.5° visual angle location had been coded NaN (i.e., was missing), the fixation duration of the corresponding time point was set to NaN and ignored during averaging.

Spatial gaze dispersion is a measure of the general tendency for the eyes to move around. It was calculated as the standard deviation in gaze, averaged across x- and y-coordinates, and transformed to logarithmic values. Smaller values indicate less gaze dispersion. To obtain time courses for gaze dispersion, it was calculated for 1-s sliding time windows centered sequentially on each time point. If less than 10% of data were available within a 1-s time window (that is, fewer than 50 samples were not NaN-coded), gaze dispersion for the corresponding time point was set to NaN and ignored during averaging.

Fixation duration and gaze dispersion do not make any assumptions about the type of eye movements under investigation and may thus be uniquely sensitive to individuals listening to masked speech. Nevertheless, we also analyzed saccade/microsaccade rate using a method that computes thresholds based on velocity statistics from x- and y-coordinate trial time courses (NaN-coded data were interpolated) and then identifies saccades/microsaccades as events passing that threshold (Engbert and Kliegl, 2003; Engbert, 2006). That is, the vertical and horizontal eye movement time series were transformed into velocities, separately for each trial, and saccades/mircosaccades were classified as outliers if they exceeded a relative velocity threshold of several times the standard deviation of the eye-movement velocity and persisted for 6 ms or longer. Previous work differed in the specific threshold that was used. Some works used a velocity threshold of 5 (Engbert and Kliegl, 2003; Widmann et al., 2014), whereas others used a threshold of 15 (Kadem et al., 2020). A lower threshold leads to a higher number of data points that are considered a saccade/microsaccade. It is unclear whether the threshold may affect the sensitivity to speech masking. Hence, we calculated analyses for both thresholds. A time course of saccade/microsaccade rate was calculated from the individual saccade/microsaccade times (Widmann et al., 2014; Kadem et al., 2020) by convolving each occurrence with a Gaussian function (standard deviation of 0.02 s; zero phase lag). We do not distinguish between saccades and microsaccades, because the definition depends on the velocity threshold and previous work suggests that the same mechanisms underlie saccades and microsaccades (Martinez-Conde et al., 2009; Martinez-Conde et al., 2013).

### Statistical analysis

The experiments reported here were not preregistered. Experimental manipulations were within-participants factors. Differences between experimental conditions were thus assessed using one-sample t-tests, paired-samples t-tests, and repeated-measures analyses of variance (rmANOVAs). Reporting of statistical results includes test statistic, degrees of freedom, significance level, and effect size. Details about statistical analyses are provided for each experiment separately below. Effect sizes for rmANOVAs and t-tests are reported as omega squared (ω^2^) and Cohen’s d (d), respectively. All statistical analyses described were carried out using MATLAB (MathWorks) and JASP (version 0.16.34) software.

## Experiment 1: The influence of speech masking on eye movements

Central fixation is common in pupillometry studies (Kuchinsky et al., 2013; Ohlenforst et al., 2017; Zekveld et al., 2018; Farahani et al., 2020; Kadem et al., 2020; Winn and Teece, 2021) but the restricted gaze may hinder examining the sensitivity of eye movements to listening effort. Moreover, forced fixation reflects a second task that may draw resources from speech comprehension. Experiment 1 explores the impact of fixation relative to free viewing on speech comprehension and the degree to which eye movements index speech-comprehension difficulties.

### Methods and materials

#### Participants

Twenty-six adults (median age: 23.5 years; age range: 18–30 years; 19 female, 7 male) participated in Experiment 1. Data from two additional participants were excluded, because not all blocks were recorded due to technical issues. For four out of the 26 participants, the quality of the eye-tracking and pupil data was low. That is, over 30% of trials contained more than 40% of missing data (Kadem et al., 2020). Hence, data from twenty-two adults (median age: 23 years; age range: 18–30 years; 16 female, 6 male) were available for analyses of eye movements (N=26 for behavioral analysis). Seven of the 22 participants were native English speakers, the other 15 participants were highly proficient non-native English speakers. Because our manipulations were all within-participants factors, the participants’ language status does not confound our speech-masking investigation.

#### Stimulus materials and procedure

Participants listened to short sentences spoken by a female native English speaker (mean duration: 2.5 s; range of durations: 2–3.2 s). Sentences were embedded in a 7-second 12-talker babble noise (Bilger, 1984). A sentence started 3 s after babble onset (Zhang et al., 2022). Sentences were presented either at −2 dB, +3 dB, or +8 dB SNR. Different SNRs were achieved by adjusting the sound level of the sentence, while keeping the babble level constant across trials (Ohlenforst et al., 2017; Kadem et al., 2020). This approach ensured that stimuli with different SNRs did not differ prior to sentence onset (Ohlenforst et al., 2017; Kadem et al., 2020). Assignment of SNR levels to specific sentences was randomized across participants. After each stimulus, a probe word occurred on the screen. Participants were asked to indicate whether the probe word is semantically related or unrelated to the sentence (Rodd et al., 2010a; Rodd et al., 2010b; Kadem et al., 2020). At the same time as participants were hearing the auditory stimuli, they were either presented with a blank gray screen (‘free viewing’) or with a fixation square centered on the gray screen (‘fixation’; the onset of the fixation square coincided with the onset of the babble and stayed on the screen until babble offset).

Participants were presented with six blocks of stimulation. In three blocks, participants were presented with the blank screen (free viewing), whereas in the other three blocks, participants were presented with the fixation square. Blocks with the blank screen versus fixation square alternated, and the starting block type was counterbalanced across participants. In each block, participants listened to stimuli in 8 trials per SNR level, presented in pseudorandomized order, such that a maximum of three trials of the same SNR level could occur in a row. Hence, across the experiment, participants listened to 24 trials for each SNR level and each fixation condition.

In order to examine whether participants found free viewing less exhausting than fixating across a block of stimulation, participants rated their mental workload for six statements after each block. Statements were ‘I felt this block went by slowly.’, ‘I found this block exhausting.’, ‘I feel tired of listening.’, ‘I had to invest a lot of effort during this block.’, ‘I felt the block was mentally demanding.’, ‘I look forward to a short break.’. Participants rated each of these statements using a 7-point scale where 1 referred to ‘strongly disagree’ and 7 referred to ‘strongly agree’. No difference in the averaged ratings was found between fixation and free viewing (t_25_ = 0.977, p = 0.338, d = 0.192).

#### Analysis of behavioral data

The proportion of correct responses in the semantic-relatedness task was calculated for each SNR condition. A rmANOVA was calculated using the proportion of correct responses as dependent measure. Within-participants factors were SNR (−2, +3, +8 dB SNR) and Viewing Condition (free viewing, fixation).

#### Analysis of eye-movement and pupil-area data

Continuous pupil-area, x-coordinate, and y-coordinate data were divided into single-trial time courses ranging from −1 to 7 s time-locked to babble onset. Data for an entire trial were excluded from analysis if the percentage of NaN data entries made up more than 40% of the trial (Kadem et al., 2020).

Fixation duration, gaze dispersion, saccade/microsaccades rate, and pupil area were calculated for each trial and then averaged across trials, separately for each SNR condition. For statistical analyses, fixation duration, gaze dispersion, saccade / microsaccades rate were averaged across the 3–5.5-s time window during which sentences were presented (sentences started at 3 s after babble onset; average sentence offset was at 5.5 s). Mean pupil area was calculated for the 4–6.5-s (i.e., delayed by 1 s), because the pupil dilation is known to change relatively slowly (Knapen et al., 2016; Winn and Moore, 2018; Winn et al., 2018; Zhang et al., 2022). An rmANOVA with the within-participants factors SNR (−2, +3, +8 dB SNR) and Viewing Condition (free viewing, fixation) was calculated separately for fixation duration, gaze dispersion, saccade/microsaccade rate, and pupil area.

### Results

#### Better speech comprehension during challenging listening when eye movements are reduced

Behavioral performance in the semantic-relatedness task decreased with decreasing SNR (F_2,50_ = 57.918, p = 9.6 · 10^-14^, ω^2^ = 0.511), but this SNR-related reduction differed depending on whether individuals freely moved or fixated their eyes (SNR × Viewing Condition interaction: F_2,50_ = 4.903, p = 0.011, ω^2^ = 0.062; Figure 1). The reduction in performance from +8 dB to −2 dB SNR was greater for free viewing than fixation (t_25_ = 2.886, p = 0.008, d = 0.566), indicating that speech-comprehension measures may be less sensitive to speech masking during fixation than free viewing. Interestingly, under the most difficult listening condition (−2 dB SNR), a reduction in eye movements (fixation) was associated with better speech comprehension relative to free viewing (t_25_ = 2.186, p = 0.038, d = 0.429), which is consistent with the hypothesis that reduced movements support listening under challenges.

**Figure 1:**
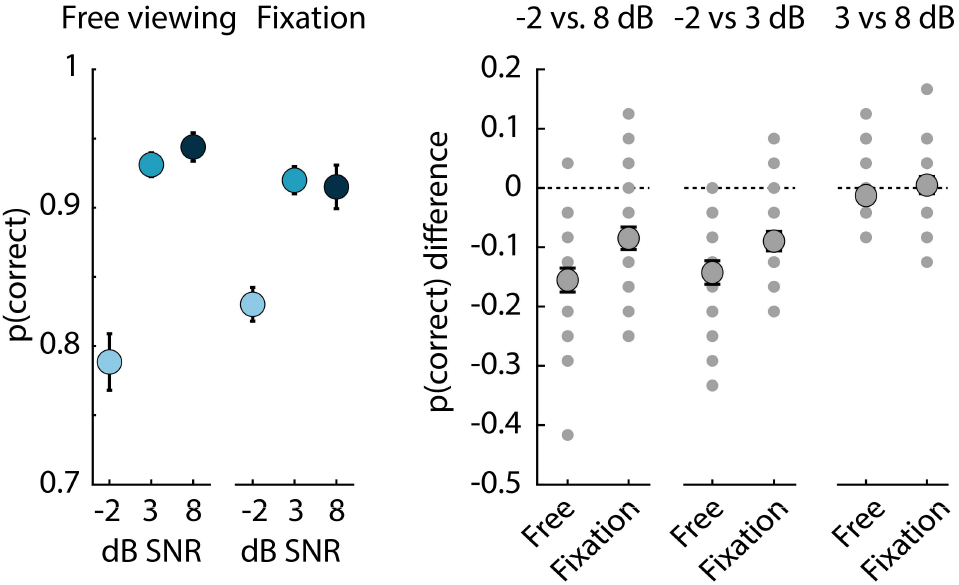
Reduced sensitive of behavioral performance to speech masking under fixation. Proportion of correct responses in the semantic-relatedness task (left) and the difference in the proportion of correct responses between different SNRs (right). The reduction in speech comprehension for −2 dB relative to 8 dB SNR was greater for free viewing than fixation (p<0.05). Speech comprehension was also greater for fixation than free viewing under challenging listening conditions (−2 dB SNR; p<0.05). Error bars reflect the standard error of the mean. Small gray dots reflect data from individual participants (some individual data points overlap in the plots).

#### Eye movements decrease and pupil area increases with increasing speech masking

Fixation durations were shorter during free viewing than during fixation, as expected (F_1,21_ = 12.069, p = 0.002, ω^2^ = 0.049). Critically, fixation durations increased with decreasing SNR (F_2,42_ = 5.752, p = 0.006, ω^2^ = 0.006; Figure 2A). Specifically, fixation durations were longer for −2 dB SNR compared to +3 dB SNR (t_21_ = 2.201, p = 0.039, d = 0.469) and +8 dB SNR (t_21_ = 3.423, p = 0.003, d = 0.730), whereas fixation durations did not differ between +3 and +8 dB SNR (t_21_ = 0.884, p = 0.387, d = 0.188). There was no SNR × Viewing Condition interaction (F_2,42_ = 0.056, p = 0.946, ω^2^ < 0.001).

**Figure 2:**
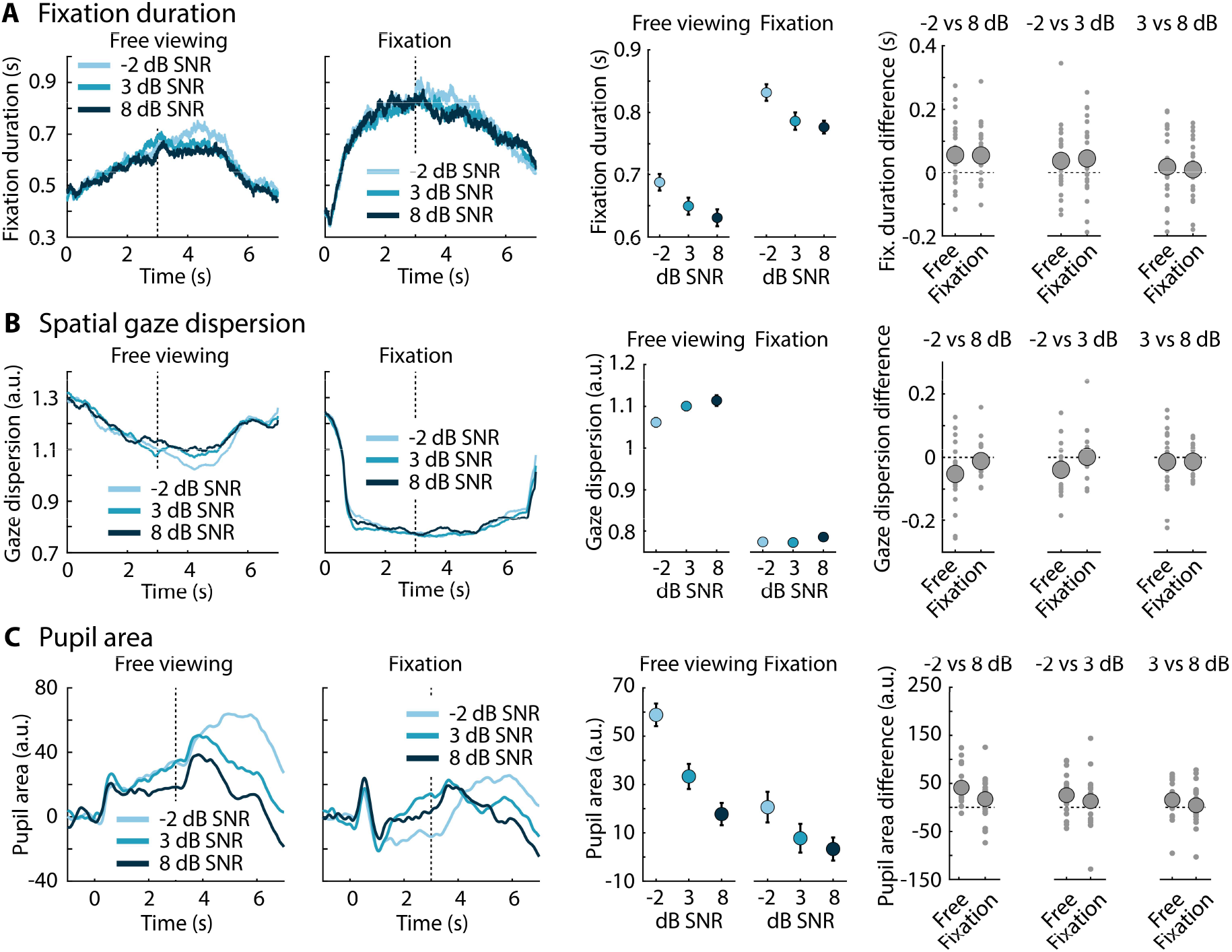
Fixation duration, spatial gaze dispersion, and pupil area are sensitive to speech masking. **A:** Left: Fixation duration time courses while participants freely viewed a blank computer screen or fixated on a fixation square at the center of the screen. The vertical dashed line in the time-course plots indicates the sentence onset. Middle: Mean fixation duration (across the 3–5.5 s time window; sentence onset at 3 s and average sentence offset at 5.5 s). Error bars reflect the standard error of the mean (removal of between-participant variance; Masson and Loftus, 2003). Right: Mean difference in fixation duration between different SNR levels for each viewing condition (free viewing, fixation). Small dots reflect data from individual participants. **B:** Same as in panel A for spatial gaze dispersion. **C:** Same as in panel A for spatial pupil area (mean across the 4–6.5 s time window).

The results for gaze dispersion were consistent with those for fixation duration. Gaze dispersion was lower during fixation than free viewing, as expected (F_1,21_ = 53.229, p = 3.5 · 10^-7^, ω^2^ = 0.213). Gaze dispersion decreased with decreasing SNR, although this effect was only marginally significant (F_2,42_ = 3.208, p = 0.051, ω^2^ = 0.001; Figure 2B). Gaze dispersion was smaller for −2 dB SNR compared to +8 dB SNR (t_21_ = 2.620, p = 0.016, d = 0.559), whereas the other contrasts were not significant (ps > 0.16). The SNR × Viewing Condition interaction was marginally significant (F_2,42_ = 2.841, p = 0.070, ω^2^ < 0.001), because gaze dispersion was only affected by SNR during free viewing (F_2,42_ = 4.050, p = 0.025, ω^2^ = 0.003), but not during fixation (F_2,42_ = 0.684, p = 0.510, ω^2^ < 0.001).

As expected based on previous work (Zekveld et al., 2010; Zekveld and Kramer, 2014; Winn et al., 2015; Winn, 2016; Winn et al., 2018; Zekveld et al., 2018; Kadem et al., 2020), pupil area increased with increasing speech masking. Pupil area was smaller during fixation than free viewing (main effect of Viewing Condition: F_1,21_ = 8.967, p = 0.007, ω^2^ = 0.037). Pupil area increased with decreasing SNR (main effect of SNR: F_2,42_ = 11.680, p = 9.3 · 10^-5^, ω^2^ = 0.035), which was due to a larger pupil area for −2 dB SNR compared to +3 dB SNR (t_21_ = 3.034, p = 0.006, d = 0.647) and +8 dB SNR (t_21_ = 4.487, p = 2 · 10^-4^, d = 0.957). Pupil area did not differ between +3 and +8 dB SNR (t_21_ = 1.806, p = 0.085, d = 0.385). The SNR × Viewing Condition interaction was not significant (F_2,42_ = 1.530, p = 0.228, ω^2^ = 0.002).

We also investigated whether the reduction in eye movements is specifically related to saccadic/microsaccadic eye movements (Figure 3). Saccade/microsaccade rate decreased with increasing speech masking, but only for a high and not the low velocity threshold defining a saccade/microsaccade. Saccade/microsaccade rate did not differ between free viewing and fixation, suggesting the measure may not be very sensitive. Specifically, the rmANOVA for saccade/microsaccade rate calculate using a threshold of 5 revealed no effect of Viewing Condition (F_1,21_ = 0.012, p = 0.913, ω^2^ < 0.001) nor of SNR (F_2,42_ = 1.098, p = 0.343, ω^2^ < 0.001) nor a SNR × Viewing Condition interaction (F_2,42_ = 2.186, p = 0.125, ω^2^ = 0.002). The rmANOVA for saccade/microsaccade rate calculated using a threshold of 15 revealed a main effect of SNR (F_2,42_ = 4.617, p = 0.015, ω^2^ = 0.007), whereas the effect of Viewing Condition (F_1,21_ = 0.038, p = 0.847, ω^2^ < 0.001) and the SNR × Viewing Condition interaction (F_2,42_ = 0.812, p = 0.451, ω^2^ < 0.001) were not significant. Saccade/microsaccade rate was lower for the −2 dB compared to the +8 dB condition (t_21_ = 3.162, p = 0.005, d = 0.674). The −2 dB versus +3 dB SNR contrast was marginally significant (t_21_ = 1.896, p = 0.072, d = 0.404), whereas no difference was found between +3 dB and +8 dB (t_21_ = 0.849, p = 0.406, d = 0.181).

**Figure 3:**
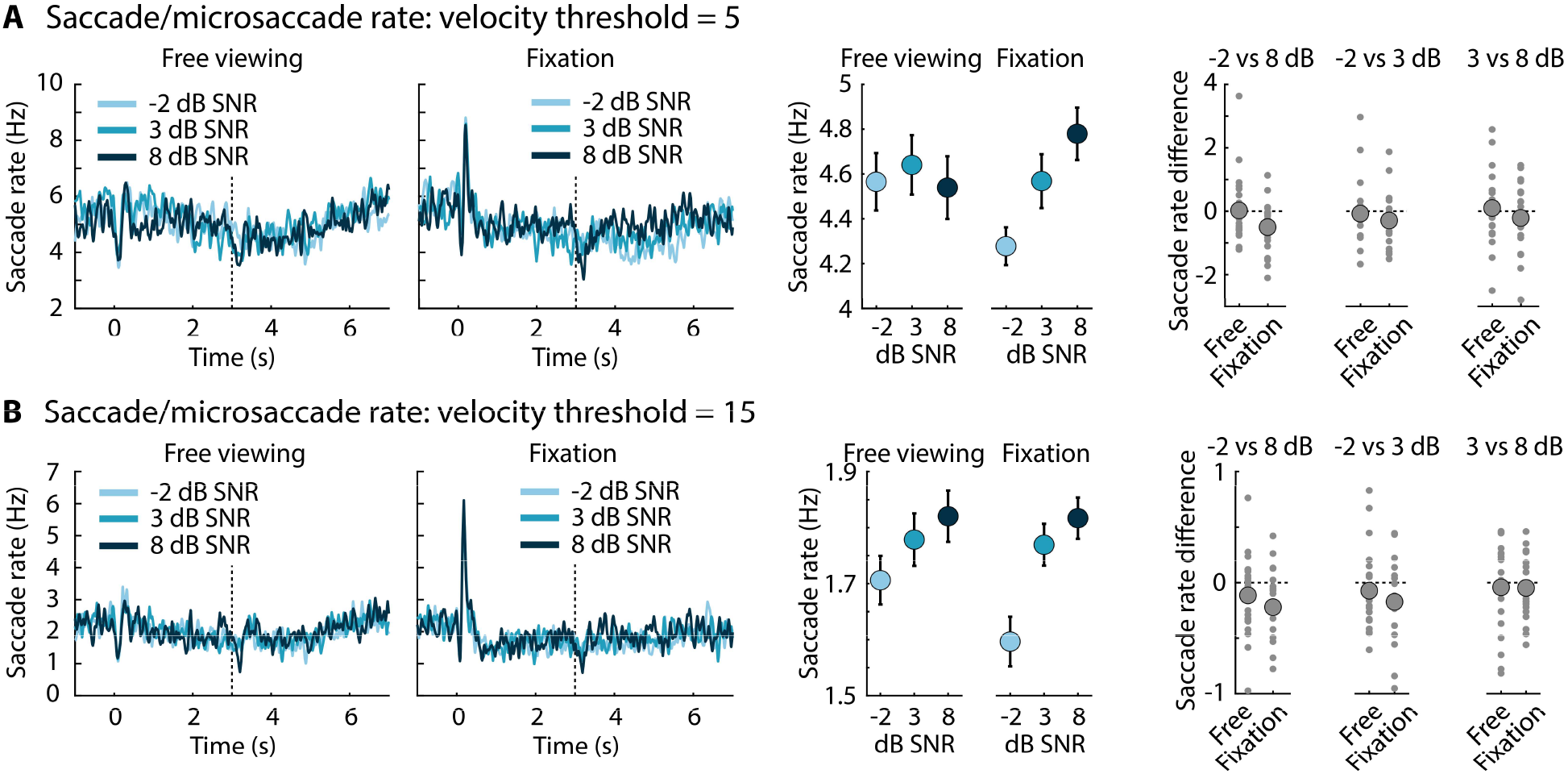
Results for saccade/microsaccade rate. **A:** Saccade/microsaccade rate time courses while participants freely viewed a blank computer screen or fixated on a fixation square at the center of the screen (left). Saccade/microsaccade rate calculated using a velocity threshold of 5. The vertical dashed line in the time-course plots indicates the sentence onset. Middle: Mean saccade/microsaccade rate (across the 3–5.5 s time window). Error bars reflect the standard error of the mean (removal of between-participant variance; Masson and Loftus, 2003). Right: Mean difference in saccade/microsaccade rate between different SNR levels for each viewing condition (free viewing, fixation). Small dots reflect data from individual participants. **B:** Same as in panel A for saccade/microsaccade rate calculated using a velocity threshold of 15.

We further investigated whether SNR-related changes in eye movements and changes in pupil area may be driven by the same underlying mechanism. To this end, we subtracted the +8 dB condition from the −2 dB condition (averaged across viewing conditions [free, fixation]), separately for pupil area, fixation duration, and spatial gaze dispersion. We calculated Pearson correlations between the difference in pupil area and the difference in fixation duration and gaze dispersion, but no significant correlation was found (fixation duration: r = 0.103, p = 0.647; gaze dispersion: r = −0.256, p = 0.250), perhaps suggesting somewhat independent processes.

Finally, the SNR effect (−2 dB minus +8 dB SNR) for behavioral performance did not correlate with the SNR effect for pupil area, fixation duration, nor gaze dispersion (fixation duration: r = −0.005, p = 0.822; gaze dispersion: r = −0.202, p = 0.368; pupil area: r = 0.377, p = 0.084), suggesting effort measures and behavioral performance dissociate to some extent (in line with Koelewijn et al., 2018; Carolan et al., 2022).

In sum, Experiment 1 shows that eye movements decrease when listening is effortful due to speech masking (Figure 2) and that speech comprehension under challenging listening is better when individuals reduce their eye movements (Figure 1). Our data also show that forced fixation reduces the sensitivity of behavioral measures of speech comprehension relative to free viewing (Figure 1). The results thus suggest that eye movements provide a window onto the perceptual/cognitive load during masked speech listening. The current results further suggest that eye movements could potentially be used to assess listening effort under non-fixation conditions and conditions that involve visual exploration. We conducted Experiment 2 to test directly whether eye movements are also sensitive to speech masking when individuals can engage in visual exploration.

## Experiment 2: The influence of speech masking on eye movements during incidental object tracking

### Methods and materials

#### Participants

Twenty-two adults (median age: 23 years; age range: 18–32 years; 14 female, 7 male; 1 person did not provide demographic information but was recruited from the same participant pool) participated in Experiment 2. Data from one additional participant were recorded but excluded from analysis, because the person’s behavioral performance was at chance level even for the easy speech-comprehension condition. Fifteen of the 22 participants were native English speakers, the other 6 participants were highly proficient non-native English speakers (the one person who did not provide demographic information was a highly proficient, likely native, English speaker).

#### Stimulus materials and procedure

Recordings of sentences from the Harvard sentence lists (mean duration: 2.4 s; range of durations: 1.9–2.9 s) (IEEE, 1969) spoken by a male native English speaker were used in Experiment 2. Sentences were embedded in a 6-second 12-talker babble noise (Bilger, 1984). A sentence started 2.5 s after babble onset. Sentences were presented either at −3 dB SNR or at +10 dB SNR. As for Experiment 1, different SNRs were achieved by adjusting the sound level of the sentence, while keeping the babble level constant across trials (Ohlenforst et al., 2017; Kadem et al., 2020). Assignment of SNR levels to specific sentences was randomized across participants. After each stimulus, a probe word occurred on the screen, and participants performed the semantic-relatedness judgement task (Rodd et al., 2010a; Rodd et al., 2010b; Kadem et al., 2020).

At the same time as participants were hearing the auditory stimuli, they were presented with incidental object-viewing stimulation (Figure 4A). Prior to babble onset, a dot was presented at the center of the screen for 0.5 s at a supra-threshold background-to-dot contrast. The dot during this 0.5-s period was colored yellow or green depending on the sentence-to-babble SNR. That is, the color of the dot served as a cue to uniquely indicate the sentence-comprehension difficulty (100% valid cue). The assignment of the colors yellow and green to the SNR levels −3 dB and +10 dB was counterbalanced across participants. Many participants reported not using the color cue; the color differences may have been too subtle. Upon babble onset, the dot turned to a light gray and started to move in a random, smooth trajectory for the 6-s duration of a trial (within a 14° visual angle; Figure 4A). Participants were instructed that the task was to comprehend the sentence so that they would be able to decide whether the probe word was semantically related or unrelated to the sentence. No task was associated with the object-movement display. Participants were instead instructed to look at the screen in whatever way they wanted (Johansson et al., 2006; Johansson et al., 2011; Johansson et al., 2012).

**Figure 4:**
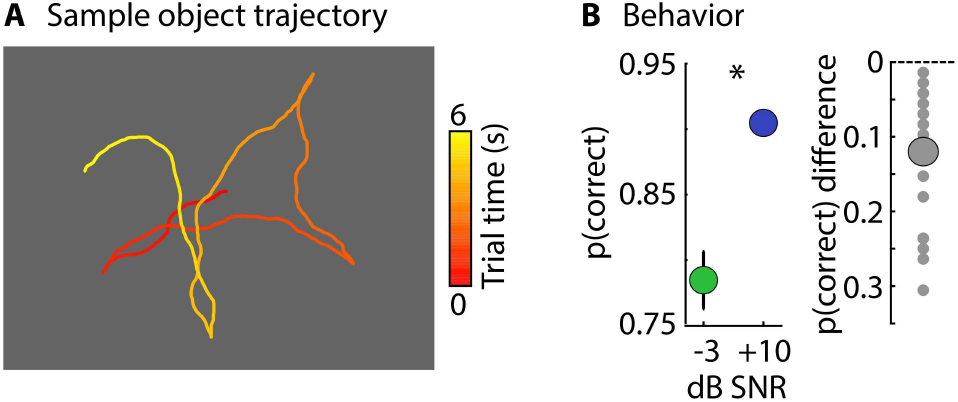
Stimulation and behavioral results for Experiment 2. **A:** Sample of an object-movement trajectory over the 6-s duration of a trial. Color indicates the temporal evolution of the movement trajectory. **B:** Left: Proportion of correct responses in the semantic-relatedness task. Error bars reflect the standard error of the mean. Right: Mean difference in the proportion of correct responses between the two SNR levels (−3 dB SNR minus +10 dB SNR). Small dots reflect data from individual participants.

Participants were presented with four blocks of stimulation. In each block, participants listened to 18 trials of each of the two SNR conditions presented in pseudo-randomized order such that a maximum of three trials of the same SNR condition were presented in a row. Hence, across the experiment, participants listened to 72 trials per SNR condition (18 trials × 4 blocks).

#### Analysis of behavioral data

The proportion of correct responses in the semantic-relatedness task was calculated for each SNR condition. A dependent-samples t-test was used to compare the proportion of correct responses between the −3 dB and the +10 dB SNR condition.

#### Analysis of eye-movement and pupil-area data

Continuous pupil-area, x-coordinate, and y-coordinate data were divided into single-trial time courses ranging from −1 to 6 s time-locked to babble onset. Data for an entire trial were excluded from analysis if the percentage of NaN data entries made up more than 40% of the trial (Kadem et al., 2020).

Fixation duration was calculated as described above. Spatial gaze dispersion was calculated slightly differently in Experiment 2 than for Experiment 1 to account for variance associated with the movement of the object. The standard deviation in gaze (averaged across x and y coordinates) was calculated and divided by the standard deviation of the object movement (averaged across x and y coordinates), and transformed to logarithmic values. Results for the normalized and the non-normalized gaze dispersion (calculated as for Experiment 1) were qualitatively similar. In addition to fixation duration and gaze dispersion, we also calculated Pearson’s correlation between the object’s and the eye’s x- and y-coordinates, and transformed the mean result to Fisher’s Z scores. To obtain time courses of the eye metrics, fixation duration was calculated for each time point, and gaze dispersion and object tracking were calculated for 1-second sliding time windows centered sequentially on each time point.

Time courses for fixation duration, gaze dispersion, Fisher’s Z scores (object tracking), saccade/microsaccade rate, and pupil area were averaged across trials. For statistical analyses, fixation duration, gaze dispersion, Fisher’s Z scores, and saccade/microsaccade rate were averaged across the 2.5– 4.9-s time window during which sentences were presented (sentences started at 2.5 s after babble onset; average sentence offset was at 4.9 s). For the analysis of the pupil area, the mean pupil area in the 3.5–5.9 s time window was calculated (again shifted by 1 s to account for delayed responsivity of the pupil; Knapen et al., 2016; Winn and Moore, 2018; Winn et al., 2018; Zhang et al., 2022). A dependent-samples t-test was calculated to compare fixation duration, gaze dispersion, Fisher’s Z scores, saccade/microsaccade rate and pupil area between the two SNR levels (−3, +10 dB SNR).

### Results

Behavioral data are displayed in Figure 4B. The proportion of correct responses in the semantic-relatedness task was lower for the −3 dB SNR compared to the +10 dB SNR condition (t_21_ = 6.494, p = 2 · 10^-6^, d = 1.385), as expected.

Analyses for eye movements showed that fixation duration increased (t_21_ = 2.775, p = 0.011, d = 0.592), and both gaze dispersion (t_21_ = 2.824, p = 0.010, d = 0.602) and object tracking decreased (t_21_ = 2.798, p = 0.011, d = 0.597) for the more challenging (−3 dB SNR) compared to the more favorable SNR (+10 dB SNR; Figure 5A-C). These data show that eye movements decrease when speech masking increases and makes listening effortful during incidental object tracking. Pupil area was larger for speech presented at −3 dB compared to +10 dB SNR, but this was only marginally significant (t_21_ = 1.794, p = 0.087, d = 0.382; Figure 5D).

**Figure 5:**
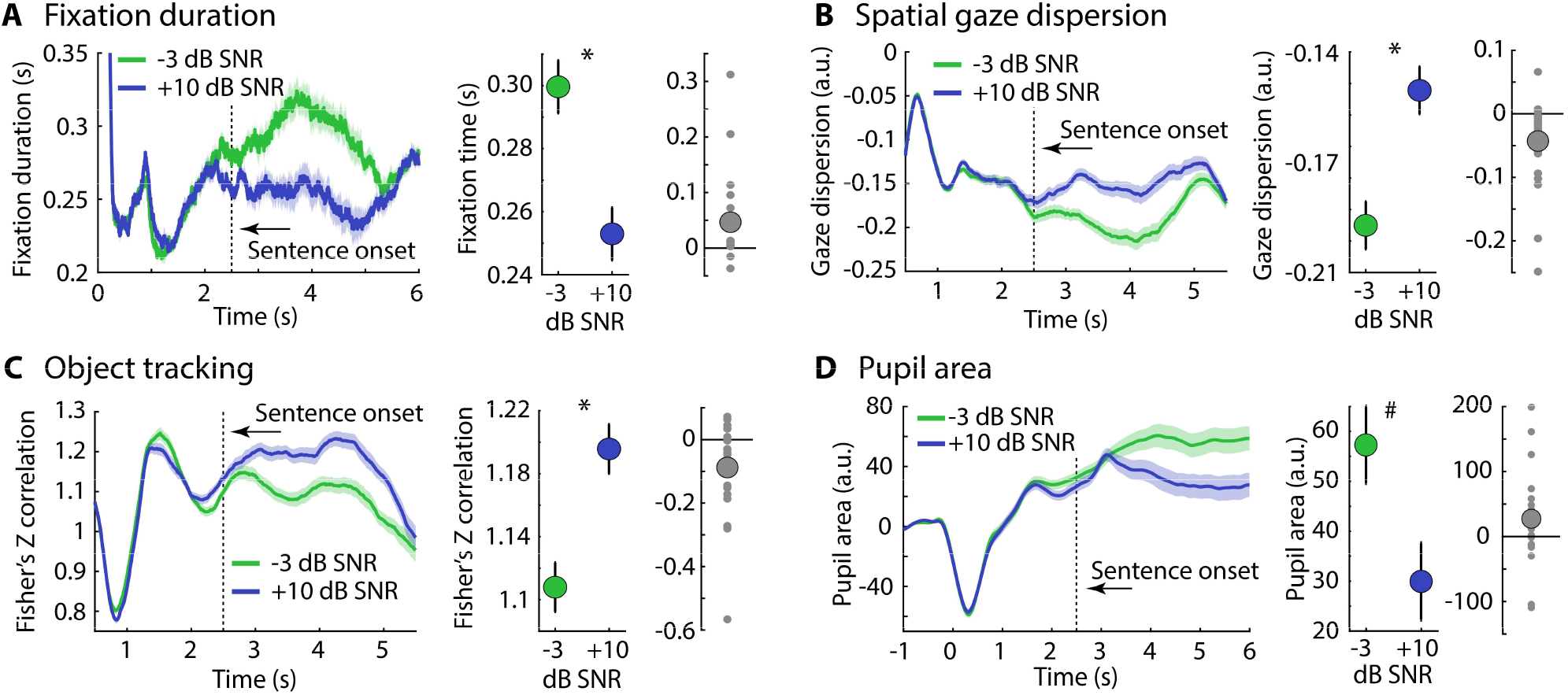
Fixation duration, gaze dispersion, object tracking, and pupil area are sensitive to speech masking: **A:** Fixation duration time courses (left). The vertical dashed line marks the sentence onset. Middle: Mean fixation duration (across the 2.5–4.9 s time window; sentence onset at 2.5 s and average sentence offset at 4.9 s). Error bars and shading reflect the standard error of the mean (removal of between-participant variance; Masson and Loftus, 2003). Right: Mean difference in fixation duration between the two SNR levels (−3 dB SNR minus +10 dB SNR). Small dots reflect data from individual participants. **B:** Same as in Panel A for spatial gaze dispersion. **C:** Same as in Panel A for Fisher’s Z transformed correlation between object and eye position (object tracking). **D:** Same as in Panel A for pupil area (mean across the 3.5–5.9 s time window). *p ≤ 0.05, #p ≤ 0.1

Saccade/microsaccade rate was lower for −3 dB compared to +10 dB SNR, but only for a low velocity threshold that defines a saccade/microsaccade (t_21_ = 2.374, p = 0.027, d = 0.506; Figure 6A) and not for the high velocity threshold (t_21_ = 1.497, p = 0.149, d = 0.319; Figure 6B), again indicating the measure may not be very sensitive (in fact, in Experiment 1, the SNR effect was only significant for the high velocity threshold).

**Figure 6:**
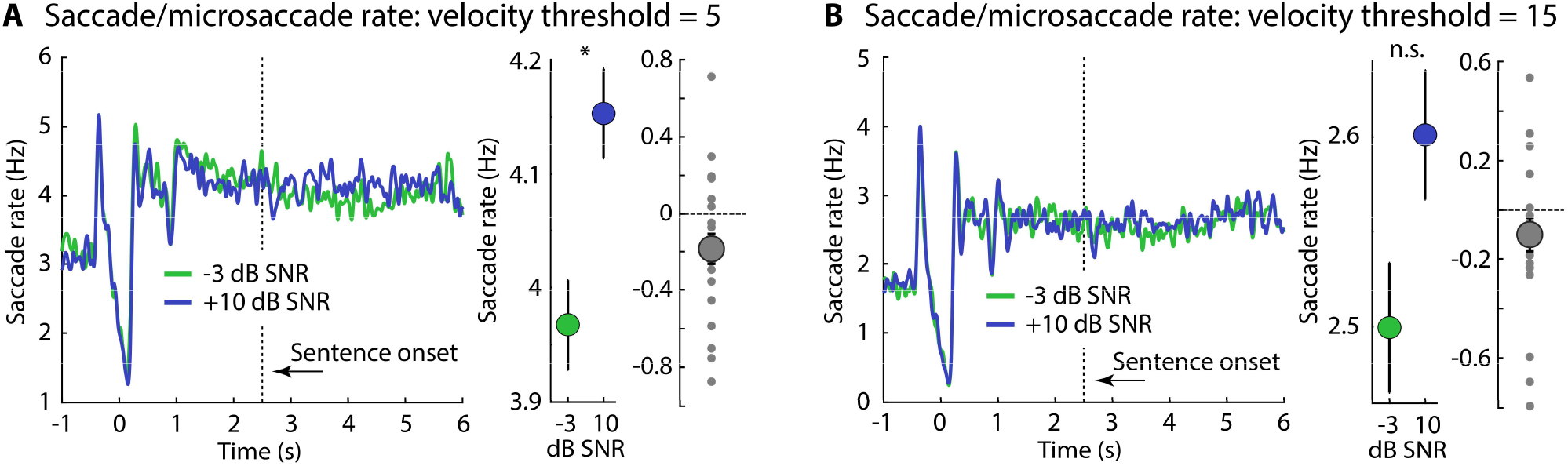
Results for saccade/microsaccade rate: **A:** Saccade/microsaccade rate time courses for velocity threshold of 5 (left). The vertical dashed line marks the sentence onset. Middle: Mean saccade/microsaccade rate (across the 2.5–4.9 s time window; sentence onset at 2.5 s and average sentence offset at 4.9 s). Error bars reflect the standard error of the mean (removal of between-participant variance; Masson and Loftus, 2003). Right: Mean difference in fixation duration between the two SNR levels (−3 dB SNR minus +10 dB SNR). Small dots reflect data from individual participants. **B:** Same as in panel A for saccade/microsaccade rate calculated using a velocity threshold of 15. *p ≤ 0.05; n.s. – not significant

Similar to observations in Experiment 1, SNR-related changes (−3 dB minus +10 dB SNR) in pupil area did not correlate with changes in eye-movement metrics (fixation duration: r = 0.255, p = 0.253; gaze dispersion: r = −0.287, p = 0.253; object tracking: r = −0.374, p = 0.087) nor did the SNR effect in behavioral performance correlate with eye-movement metrics or pupil area (fixation duration: r = −0.066, p = 0.771; gaze dispersion: r = 0.02, p = 0.93; object tracking: r = 0.005, p = 0.984; pupil area: r = 0.182, p = 0.418).

In sum, Experiment 2 expands the results of Experiment 1, showing that eye movements decrease when speech masking increases under incidental object tracking. Pupil dilation appears slightly less sensitive to speech-comprehension difficulties than eye-movement metrics in Experiment 2. The results of Experiment 1 and 2 provide a clear demonstration that eye movements carry critical information about the challenges during speech comprehension and that this sensitivity of eye movements to masked speech listening generalizes to different visual-stimulation conditions. Importantly, listening situations not only vary in the visual information available, but also in the type of speech materials. Experiment 3 examines whether eye movements are also sensitive to listening effort for engaging, continuous speech materials that are common in everyday life (Jefferson, 1978; Mullen and Yi, 1995; Bohanek et al., 2009; Fivush et al., 2011).

## Experiment 3: The influence of speech masking on eye movements during story listening

Difficulties with speech comprehension and associated effort have been assessed mostly using brief, disconnected sentences (Zekveld et al., 2010; Wendt et al., 2016; Ayasse and Wingfield, 2018; Zekveld et al., 2019; Kadem et al., 2020; Winn and Teece, 2021; cf. Experiments 1 and 2). Such materials lack a topical thread and are not very interesting to a listener. Speech in everyday life is often continuous, follows an overarching theme, and a listener is intrinsically motivated to comprehend (Jefferson, 1978; Mullen and Yi, 1995; Bohanek et al., 2009; Fivush et al., 2011; Herrmann and Johnsrude, 2020b). Any measure of listening effort would ideally be sensitive to such continuous, naturalistic speech. However, pupillometry may be less suited for the assessment of listening effort during continuous speech listening, because measures of pupil dilation typically require normalization to a pre-sentence baseline period (Zekveld et al., 2010; Winn et al., 2018; Zekveld et al., 2018; Zekveld et al., 2019; Kadem et al., 2020; Winn and Teece, 2021), which is less possible for continuous speech. The eye-movement metrics established in Experiments 1 and 2 (i.e., fixation time and spatial gaze dispersion) do not require baseline normalization and may thus be uniquely sensitive to listening effort during continuous speech listening.

### Methods and materials

#### Participants

Twenty-three adults (median age: 25 years; age range: 18–34 years; 15 female, 6 male, 1 non-binary, 1 person did not answer) participated in Experiment 3. Data from one additional participant were recorded but excluded from analysis, because more than 40% of their data were rejected during preprocessing. Twenty-one of the 23 participants were native English speakers, the other 2 participants were highly proficient non-native English speakers.

#### Acoustic stimulation and procedure

Participants listened to two approximately 10-min stories from the storytelling podcast The Moth (https://themoth.org/; “Nacho Challenge” by Omar Qureshi [11 min]; “Family trees can be dangerous” by Paul Nurse [10 min]). The Moth is a podcast where people tell stories about interesting life events. Stories are highly enjoyable and absorbing (Herrmann and Johnsrude, 2020a; Irsik et al., 2022a). Each participant listened to one of the stories in the original temporal order (referred to as ‘intact’), whereas they listened to the other story in scrambled order, for which the order of phrases/sentences was shuffled (referred to as ‘scrambled’). A scrambled story was included as a control story that resembles short, disconnected sentences for which we expected story comprehension, listening motivation, and effort investment to be low. Scrambled stories were obtained using a custom MATLAB script as follows. Silence periods with a duration of at least 0.05 s were identified. The duration of speech snippets separated by the identified silence periods was calculated. Speech snippets with a duration of 1 s or longer were cut out at the center of the silence periods and the order of snippets was shuffled. Story scrambling was calculated for each participant uniquely. Which of the two The Moth stories was ‘intact’ versus ‘scrambled’ was counterbalanced across participants. The order in which the ‘intact’ versus the ‘scrambled’ story was presented was also counterbalanced across participants.

Each story was masked by 12-talker background babble (Bilger, 1984). The SNR between the speech signal and the 12-talker babble changed every 28 seconds to one of five SNR levels (+16, +11, +6, +1, −4 dB SNR; for a similar approach see Irsik et al., 2022a, b), corresponding to about a range of 95% to 50% of intelligible words (Irsik et al., 2022a). The SNR was manipulated by adjusting the dB level of both the story and the masker. This ensured that the overall sound level remained constant throughout a story and was similar for both stories. Each story started and ended with the +16-dB SNR level to enable participants to clearly hear the beginning and end of the story. Each SNR level was presented 4 times, with the exception of the +16 dB SNR level, which was presented 6 times (4 times + beginning and end). The SNR transitioned smoothly from one level to the other over a duration of 1 second. The order of SNR levels was randomized such that a particular SNR could not be heard twice in succession, and that SNR would maximally change by two levels. For each participant, SNR levels were randomized uniquely, but the same randomization was used for the ‘intact’ and the ‘scrambled’ story.

After a story ended, participants answered ten comprehension questions about the story. For each comprehension question, participants were asked to select the correct answer out of four multiple choice options.

#### Visual stimulation

In order to facilitate eye movements, we adopted an incidental multi-object movement display (Cavanagh and Alvarez, 2005; Alvarez and Franconeri, 2007; Scholl, 2009; Herrmann and Johnsrude, 2018b, a) that was concurrently presented with the spoken stories. To this end, 16 dots [dot diameter: 1.2 cm (0.9°)] were presented and moved on the screen. Presentation of dots was constrained to a display frame of 20.6 cm width (15.6°) and 19.4 cm height (14.7°) centered on the screen and highlighted to the participants by a black frame on a gray background (Figure 7B). Dots never moved outside of the display frame and never overlapped during movements; dots moved ~3.7 cm/s (2.8°/s). The locations of the 16 dots were set to new, randomly selected locations every 3–5 seconds to facilitate eye movements and overcome the technical challenge that displaying continuous dot movements for the ~10 min story duration exceeded the working memory capacity of the stimulation laptop.

**Figure 7:**
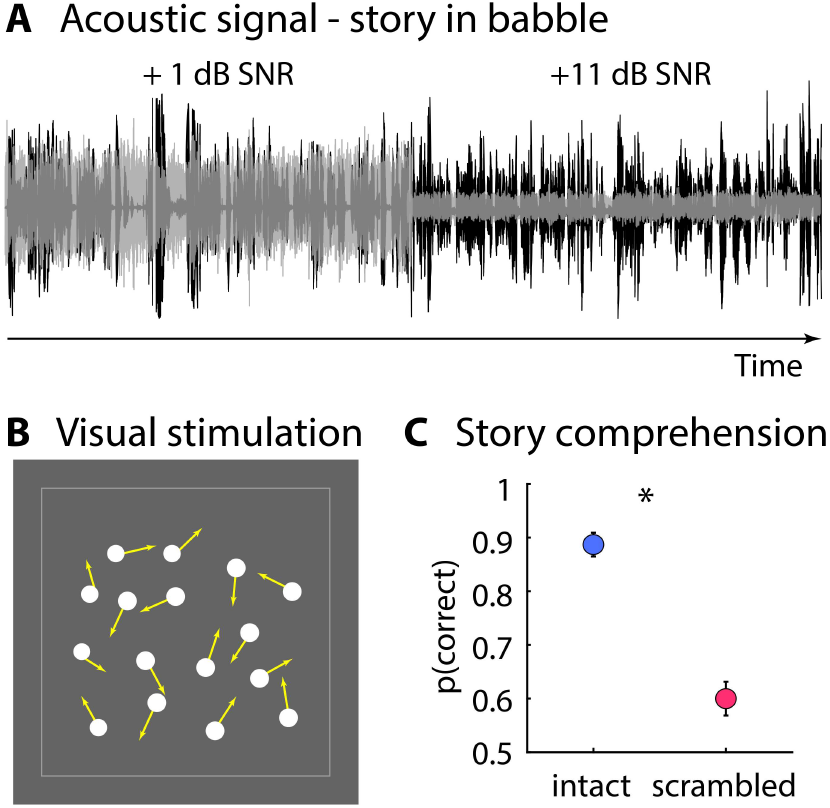
Stimulation and behavioral data for Experiment 3. **A:** Sample snippet of the engaging, spoken story masked by background babble. The signal-to-noise ratio (SNR) between the speech signal and the background babble changed every 28 seconds to one of five SNR levels (−4, +1, +6, +11, +16 dB SNR). **B:** Moving-dot display presented while participants listened to the spoken story. Sixteen dots smoothly moved on the screen. No task was required using the moving-dot display. **C:** Proportion of correctly answered story-comprehension questions for the intact and scrambled story (the temporal order of phrases/sentences were shuffled within the scrambled story). *p<0.05

Critically, participants were instructed that the task was to comprehend the story so that they would be able to answer the comprehension questions following the story presentation. Hence, participants did not need to perform a task on the multiple-object movement display. Participants were instead instructed to look at the screen in whatever way they wanted (Johansson et al., 2006; Johansson et al., 2011; Johansson et al., 2012).

#### Analysis of behavioral data

Responses to comprehension questions were coded as correct or incorrect. A mean story-comprehension score was calculated as the proportion of correct responses across the ten comprehension questions, separately for the ‘intact’ and the ‘scrambled’ story. A dependent-samples t-test was calculated to compare the proportion of correct responses between the ‘intact’ and the ‘scrambled’ story.

#### Analysis of eye-movement and pupil-area data

Fixation duration, spatial gaze dispersion, and mean pupil area were calculated for each time point over the duration of a story. Gaze dispersion and mean pupil area were calculated across a 1-s time window centered on each time point. Fixation duration, gaze dispersion, and mean pupil area were averaged across time points, separately for each of the five SNR levels and the two story types (‘intact’, ‘scrambled’). Saccade/microsaccade rate was not calculated for story-listening data, because the method to estimate what counts as a saccade/microsaccades has been developed for trial-based data (Engbert and Kliegl, 2003; Engbert, 2006; Widmann et al., 2014; Kadem et al., 2020).

For each participant and story type, a linear function was fit separately to fixation duration, gaze dispersion, and pupil area as a function of SNR levels. The resulting slope (linear coefficient) was tested against zero using a one-sample t-test in order to test whether there was a significant relation between SNR levels and the eye metrics. A dependent-samples t-test was calculated to compare the slopes between the ‘intact’ and the ‘scrambled’ story.

### Results

The proportion of correctly answered comprehension questions was greater for the intact story compared to the scrambled story, as expected (t_22_ = 8.43, p = 2.4 · 10^-8^, d = 1.758; Figure 7C).

For fixation duration, the slope of a linear function fit relating SNR to fixation duration showed that fixation duration decreased with increasing SNR for the intact (t_22_ = −2.660, p = 0.014, d = 0.555) but not for the scrambled story (t_22_ = 0.440, p = 0.664, d = 0.092; Figure 8A; difference between slopes: t_22_ = 2.043, p = 0.053, d = 0.426). In fact, mean fixation duration (across SNRs) for the scrambled story did not differ from the fixation duration for the most favorable SNR of the intact story (+16 dB; t_22_ = 0.685, p = 0.500, d = 0.143), but was smaller than the fixation duration for the most unfavorable SNR (−4 dB; t_22_ = 2.672, p = 0.014, d = 0.557). In other words, fixation duration during listening to the scrambled story mirrored the fixation duration when speech comprehension was easy during listening to the intact story, consistent with hypothesis that individuals do not invest cognitively while listening to a scrambled story.

**Figure 8:**
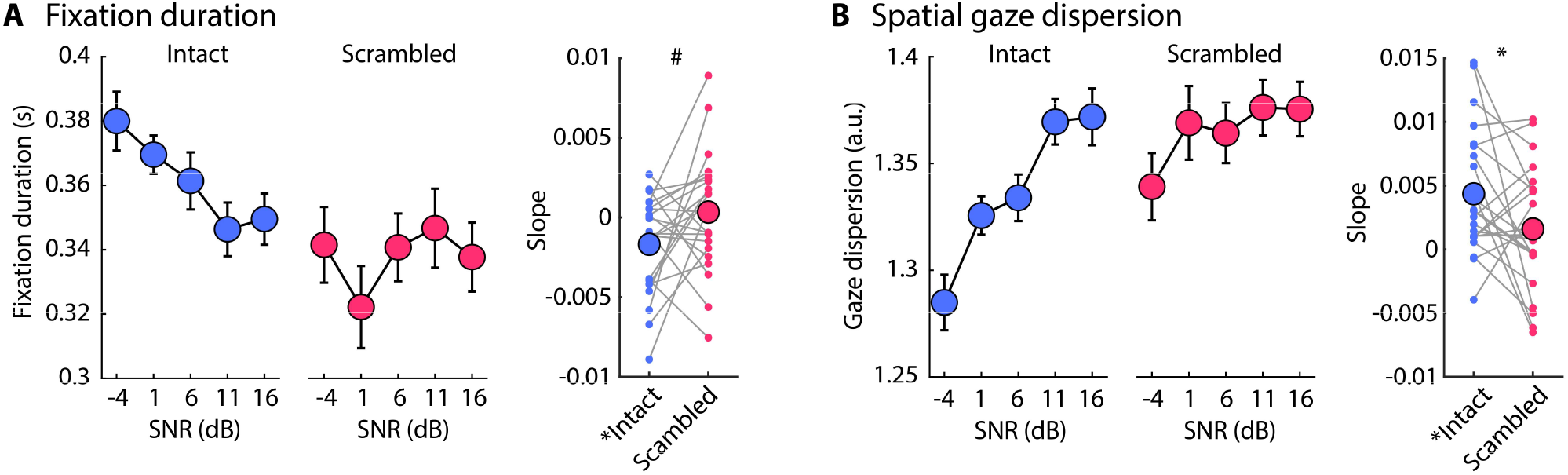
Eye movements decrease when story listening is challenging. **A:** Fixation duration for the intact and the scrambled story for each SNR level (left). The error bars reflect the standard error of the mean (removal of between-participants variance, Masson and Loftus, 2003). The right plot shows the slope from a linear function fit, reflecting the linear relation between SNR levels and fixation duration. Small dots reflect data points from individual participants. The asterisk in front of the condition label indicates a significant difference from zero (p≤0.05). **B:** Same as in panel A for spatial gaze dispersion. *p≤0.05, #p≤0.1

Results for spatial gaze dispersion mirrored those for fixation duration. Gaze dispersion increased with increasing SNR for the intact story (t_22_ = 4.250, p = 3.3 · 10^-4^, d = 0.886), but not for the scrambled story (t_22_ = 1.609, p = 0.122, d = 0.336; difference between slopes: t_22_ = 2.147, p = 0.043, d = 0.448). Moreover, mean gaze dispersion for the scrambled story (across SNRs) was not different from gaze dispersion at the most favorable SNR of the intact story (+16 dB; t_22_ = 0.257, p = 0.800, d = 0.054), but significantly higher compared to the least favorable SNR of the intact story (−4 dB; t_22_ = 3.102, p = 0.005, d = 0.647). The data thus suggest again that individuals’ eyes moved as much during the scrambled story as they did during the easiest SNR while listening to the intact story, indicating that listeners did not invest cognitively while listening to the scrambled story.

In contrast to previous and current pupil data recorded during sentence listening (Experiments 1 and 2; e.g., Zekveld et al., 2010; Wendt et al., 2016; Kadem et al., 2020), we found no linear relation between SNR and pupil area for the intact story (t_22_ = 0.053, p = 0.958, d = 0.011; Figure 9). However, there was an unexpected linear increase in pupil area with increasing SNR for the scrambled story (t_22_ = 2.163, p = 0.042, d = 0.451; Figure 9; no difference between the intact and the scrambled story: t_22_ = 1.528, p = 0.141, d = 0.319). The absence of an SNR effect on pupil area during engaging story listening, may be related to the incidental moving-dot display, reducing sensitivity, and/or due to the omission of pupil baseline normalization, which is not feasible for continuous speech listening over several minutes. The increase in pupil area with decreasing SNR perhaps reflects arousal (Bradshaw, 1967; Bradley et al., 2008; Mathôt, 2018; Ayasse and Wingfield, 2020) when speech becomes unmasked for low-engaging materials.

**Figure 9:**
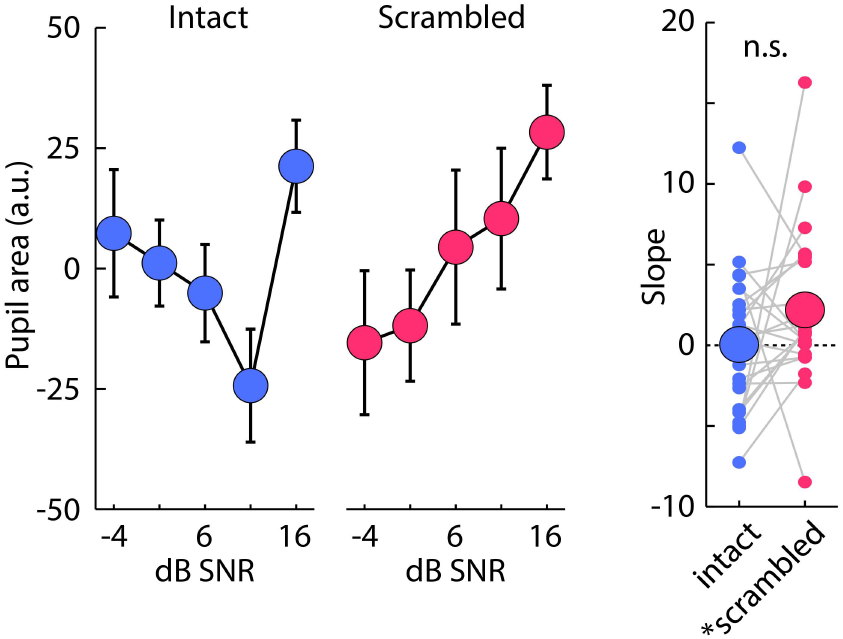
Pupil-area data. Pupil area for the intact and the scrambled story for each SNR level (left). The error bars reflect the standard error of the mean (removal of between-participants variance, Masson and Loftus, 2003). The right plot shows the slope from a linear function fit, reflecting the linear relation between SNR levels and pupil area. The asterisk in front of the condition label indicates a significant difference from zero (p≤0.05). n.s. – not significant

In sum, Experiment 3 shows that individuals’ eye movements decrease when speech is masked by background noise during listening to an engaging, continuous story. The sensitivity of eye movements to listening challenges induced by background masking was specific to the interesting, engaging story and was absent for a less engaging, scrambled story. Experiment 3 further shows that pupil dilation is less sensitive to speech masking during continuous speech listening. The current data thus highlight the challenges with pupillometric measures in more naturalistic speech-listening paradigms, and suggest that other eye-based metrics such as fixation duration and gaze dispersion may provide an alternative.

## Discussion

Pupillometry is the most used objective tool to assess listening effort but has multiple disadvantages. The current study explored a new way to assess listening effort based on eye movements. In three experiments, we show that listeners’ eye movements decrease when listening to masked speech is effortful, as indicated by increased fixation duration and decreased spatial gaze dispersion. We demonstrate this effort-related reduction in eye movements during free viewing and incidental object tracking, as well as for simple sentences and naturalistic, continuous stories. Pupillometry was not sensitive to listening effort during story listening (only during isolated sentence listening), highlighting the challenges with pupillometric measures for the assessments of listening effort in naturalistic speech-listening paradigms. Our results reveal a critical link between eye movements and cognitive load that can be leveraged to measure listening effort.

### Reduced eye movements as a measure of cognitive load

Whereas the current work is, to our knowledge, the first to demonstrate reduced eye movements during difficult listening, a few studies have shown that individuals avert gaze (Glenberg et al., 1998), reduce object-tracking eye movements (Lipton et al., 1980; Hutton and Tegally, 2005; Kosch et al., 2018), and decrease saccades (Walter and Bex, 2021) when memory load is high compared to low. It thus appears that different cognitively challenging tasks affect oculomotor function such that eye movements decrease during periods of high cognitive load.

We also observed that saccades/microsaccades are sensitive to speech masking, but this appeared to depend on the threshold that defined what counts as a saccade/microsaccade in inconsistent ways across Experiment 1 and 2 (Figures 3 and 6). Saccade/microsaccade rate was also not sensitive to the different viewing conditions (free vs fixation) in Experiment 1, whereas fixation duration and graze dispersion were. Previous research did not find that listening effort affects saccades/microsaccades, but different thresholds were not tested in this study (Kadem et al., 2020). Saccades/microsaccades occur relatively infrequently about 1 to 3 times per second (Martinez-Conde et al., 2009; Martinez-Conde et al., 2013; Pierce et al., 2019), which may make saccades/microsaccades an insensitive measure of listening effort. The eye-movement measures proposed in the current study – fixation duration and gaze dispersion – do not focus on a specific type of eye movements, but instead leverage all x- and y-data within a time window as a basis for calculations, possibly leading to increased sensitivity. Nevertheless, given the inconsistent effect of speech-masking on saccades/microsaccades, the current data may suggest that non-saccadic eye movements, possibly smooth eye movements, contribute to the reduction in eye movements during effortful speech listening.

### Different sensitivity of pupil area versus eye-movement metrics to listening effort

We show that pupil area increases with increasing effort induced by masked speech during sentence listening (Figures 2C and 5D), in line with a large body of previous work (Zekveld et al., 2010; Kuchinsky et al., 2013; Zekveld et al., 2014; Zekveld and Kramer, 2014; Winn et al., 2015; Wendt et al., 2016; Winn, 2016; Wendt et al., 2017; Zekveld et al., 2019; Kadem et al., 2020; Seifi Ala et al., 2020; Winn and Teece, 2021; Zhang et al., 2022). The current results further show that the masking-related increase in pupil area is also present when fixation on a stationary point is not required (see also Kraus et al., 2022). Fixation to a stationary point reduces the sensitivity of behavioral measures of speech comprehension (Figure 1), and may impair memory and mental imagery for spoken speech (Johansson et al., 2012). The observation that pupil area is sensitive to listening effort during free viewing (Figure 2C) and incidental object tracking (Figure 5D) perhaps suggests that strict fixation to assess listening effort with pupillometry is not required.

Pupil area was not sensitive to effort-related speech masking when individuals listened to continuous, engaging stories (Figure 9). In contrast, fixation duration increased and spatial gaze dispersion decreased with increased speech masking during both sentence and story listening (Figures 2, 5, and 8), emphasizing the potential of using eye movements to assess listening effort for naturalistic speech listening. Previous work using continuous speech materials that consisted of 30-s passages, rather than the ~2-3 s sentences typically used, observed a larger pupil size for less compared to more favorable speech-masking levels (Seifi Ala et al., 2020; Fiedler et al., 2021). Pupil size in these studies was normalized to a pre-speech baseline, which is also commonly done in sentence-listening paradigms (Winn et al., 2018; Zekveld et al., 2018). Baseline normalization is not attainable for ~10 min continuous stories because no ‘neutral’, speech-devoid time period is available. The absence of baseline normalization in our story-listening experiment may have contributed to the insensitivity of the pupil area to listening effort. The eye-movement metrics proposed here – fixation duration and spatial gaze dispersion – do not require baseline normalization, and may thus be uniquely sensitive to speech masking during story listening.

Our analyses show that changes in pupil area and changes in eye-movement metrics do not significantly correlate. Whereas the current study was not designed to specifically examine inter-individual variations (i.e., correlation analyses may require a higher number of participants; Bossier et al., 2020; Grady et al., 2021), the absence of a correlation between pupil-area and eye-movement changes may imply distinct processes or mechanisms (although note that the direction of the relation was as expected). Previous works have also failed to find a significant relationship between indices of listening effort, for example among pupil area, neural oscillation, heart rate, and skin conductance (Miles et al., 2017; Strand et al., 2018; Alhanbali et al., 2019; Seifi Ala et al., 2020; Kraus et al., 2022). The absence of a relationship between listening-effort measures might be due to different neural mechanisms that tap into different aspects of effort (Strand et al., 2018; Herrmann and Johnsrude, 2020b; Strand et al., 2020).

### Neural mechanisms of reduced eye movements during challenging listening

That eye movements reduce with increasing listening effort aligns with work in non-human mammals, showing increased activity in auditory cortex during periods of reduced movements (Schneider et al., 2014; McGinley et al., 2015; Schneider and Mooney, 2015, 2018; O’Connell et al., 2020), and heightened neuronal excitability in auditory cortex in the absence of eye movements (O’Connell et al., 2020). Reductions in any (eye) movements may thus support auditory processing, including speech perception, by enhancing auditory-system sensitivity. The decrease in eye movements may further reduce visual, proprioceptive, and other inputs that may distract cognitively from listening.

Generation and regulation of eye movements relies on a network of neural circuits that involves cortical and subcortical brain structures, such as the visual cortex, prefrontal cortex, posterior parietal cortex, frontal and supplementary eye fields, anterior cingulate cortex, cerebellum, thalamus, basal ganglia, and superior colliculus (Sparks, 2002; Pierrot-Deseilligny et al., 2004; Pierce et al., 2019). Some of these regions overlap with the network that regulates the pupil size (Wang et al., 2012; Joshi and Gold, 2020; Wang and Munoz, 2021; Burlingham et al., 2022), highlighting the potential for shared mechanisms of effort-related changes in pupil area and eye movements.

Which regions drive the reduction in eye movements during challenging listening is unclear. We speculate that cognitive control regions, such as the prefrontal, cingulate, and parietal cortices (Cole and Schneider, 2007; Braver, 2012; Niendam et al., 2012), may influence structures that initiate eye movements, such as the frontal and supplementary eye fields and the superior colliculus (Pierce et al., 2019). However, the degree of network activation has been shown to depend on whether eye movements are automatically triggered or volitionally initiated (Pierce et al., 2019). As such, the specific involvement of brain regions may depend on whether listeners consciously choose to reduce eye movements when listening becomes effortful, or whether eye movements decrease automatically. While eye movements are under cognitive control, the observation of increased fixation duration with decreasing SNR while listeners fixate on a stationary point (Figure 2) perhaps suggests involuntary contributions. The question of whether the reduction of eye movements is deliberate versus involuntary has also implications for the type of effort the reduced eye movements index. A deliberate reduction implies that the listener experiences listening effort first and subsequently reduces eye movements, whereas an involuntary process may index cognitive resource recruitment rather than the experience of effort (for important recent discussions on experienced vs exerted effort see Lemke and Besser, 2016; Herrmann and Johnsrude, 2020b; Strand et al., 2020).

## Conclusions

Pupillometry is the most used objective tool to assess listening effort but has several disadvantages. Building on cognitive and neurophysiological work, the current study explores a new, objective way to assess listening effort through eye movements. Here, we examine the hypothesis that eye movements decrease when speech listening becomes effortful. Consistent with this hypothesis, we demonstrate that fixation duration increases and spatial gaze dispersion decreases with increasing speech masking. Eye movements decrease when speech comprehension is effortful during free viewing and object tracking, as well as for sentence and story listening. Pupillometry was sensitive to speech masking only during sentence listening, but not during story listening, highlighting the challenges with pupillometric measures for the assessments of listening effort during naturalistic speech listening. Our results reveal a critical link between eye movements and cognitive load during speech comprehension, and provide the foundation for a novel measure of listening effort that has the potential to be applicable in a wide range of contexts.

## Author contributions

MEC designed research, performed research, analyzed data, and revised the paper. BH designed research, performed research, analyzed data, and wrote the paper.

## Acknowledgements

The research was supported by the Canada Research Chair program (232733), the Natural Sciences and Engineering Research Council of Canada (Discovery Grant: RGPIN-2021-02602), and the William Demant Foundation. We thank Devin Sodums for this help during data collection. We thank Ingrid Johnsrude, Jennifer Ryan, and Lauren Fink for fruitful discussions.

